# Development and Optimization of ^111^In-Dinutuximab-IRDye800, a Dual-Modality Intraoperative Molecular Imaging Agent for Pediatric Neuroblastoma Resection

**DOI:** 10.64898/2026.08.28.747876

**Authors:** Catherine Y. Yip, Lauren T. Rosenblum, Arjun Pant, Arianna Kahler-Quesada, Bhuvitha Chagantipati, ReidAnn Sever, Brady Grano-Mickelsen, Bo Li, Angel G. Cortez, Joseph D. Latoche, Kathryn E. Day, Lora Rigatti, Jessie R. Nedrow, W. Barry Edwards, Gary Kohanbash, Marcus M. Malek

## Abstract

**Rationale:** Neuroblastoma is a devastating pediatric malignancy, for which surgical resection is a key factor in long-term survival. However, there are significant challenges in its resection, particularly in high-risk disease, as neuroblastoma encases surrounding critical structures, is often difficult to distinguish from desmoplastic or scar tissue, and can carry occult deposits of disease not readily identified on preoperative imaging or intraoperative visualization. Building on the principles of fluorescent and radio-guided surgery, in combination with the known overexpression of GD2 in neuroblastoma, we sought to develop and optimize ^111^In-Dinutuximab-IRDye800, a dual-modality GD2-targeted intraoperative molecular imaging agent, for use in pediatric neuroblastoma to help enhance patient safety while facilitating a more complete resection.

**Methods:** Dinutuximab was conjugated to IRDye800 and DTPA, then radiolabeled with Indium-111 to yield ^111^In-Dinutuximab-IRDye800. Optimization occurred through ELISA assay to assess binding affinity, fluorescence intensity analysis to determine the optimal fluorescent degree of labeling, and phototoxicity testing through flow cytometry. Rodent models of neuroblastoma were then generated through injection of SK-N-BE(2) human neuroblastoma cells into the left adrenal glands of nude mice or RNU rats. A series of fluorescent and gamma biodistributions was performed, varying the dose, timing, and specific activity of the tracer. Tumor and organ uptake of the tracer was compared with one- or two-way ANOVA as appropriate, with Sidak’s multiple comparison test to compare tumor uptake to individual organs. Once optimization was complete, a clinically significant events study modeled after human clinical trials was performed to evaluate the *in vivo* capabilities of ^111^In-Dinutuximab-IRDye800.

**Results:** Increased ratios of IRDye800 per antibody led to decreased binding affinity for GD2 and was associated with formulation instability without significant return on fluorescence intensity. Specific activity of the tracer was not found to impact overall biodistribution of the tracer. A 45-50 µg dose of ^111^In-Dinutuximab-IRDye800 with ratios around 1 DTPA and 1-1.5 IRDye800 per antibody imaged 4 days after tracer administration was found to be the optimal combination that maximized detectable tumor-specific signal. In the clinically significant events study mirroring human IMI clinical trials, fluorescent guidance identified additional malignant lesions not originally detected under white light in 64% of rodents.

**Conclusions:** ^111^In-Dinutuximab-IRDye800 is a dual-modality GD2-targeted intraoperative imaging agent that is well-poised for clinical translation. As it preserves tumor specificity, yields clinically meaningful radiofluorescent signal, and is well-tolerated without adverse events after optimization was completed, it carries the potential to positively impact the safety and completeness of neuroblastoma resection.

## Introduction

Neuroblastoma is the most common extracranial solid tumor in children, accounting for 8% of all childhood cancers but a disproportionate 15% of pediatric cancer deaths [1, 2]. Although low- and intermediate-risk groups carry overall good prognoses and long-term survival, survival of high-risk neuroblastoma remains dismal, around or below 50% [3] despite intensive multimodal therapy consisting of chemotherapy, surgical resection, radiation, and immunotherapy.

As surgical resection is a key component of standard treatment regimens, studies evaluating its role on outcome have demonstrated that achieving complete macroscopic excision or ≥90% resection is associated with improved event-free survival (EFS) and decreased rates of local progression [4, 5]. However, the rate of inadequate resection in pediatric neuroblastoma can be as high as 30-50% [6]. There is potential for significant surgical morbidity as neuroblastoma typically encases critical neurovascular structures, which cannot simply be resected with the tumor. The risk of surgical complications, including significant blood loss, nerve or vessel injury, and unplanned organ resection, therefore remains high, approaching 30-50% [5]. Specific challenges in resection of high-risk neuroblastoma include distinguishing tumor from normal tissue or scar tissue that has formed as a result of neoadjuvant chemotherapy, balancing the goal of ≥90% resection with preservation of critical structures, and identification of remote deposits of disease.

Although preoperative imaging aids in surgical planning and identifying patients at risk of surgical complications, there is still a lack of real-time, intraoperative tumor-specific guidance during resection. This challenge in pediatric neuroblastoma and other malignancies has driven the field of intraoperative molecular imaging (IMI), building on the foundations established by fluorescence-guided surgery (FGS) and radio-guided surgery (RGS). Traditionally, FGS and RGS rely on passive accumulation of the tracer based on properties of the molecule and tissue itself [7]. Sentinel lymph node biopsy for breast and skin malignancies is among the most common clinical use of RGS, relying on the passive accumulation of a non-specific tracer into lymphatic channels based on molecular size [8, 9]. While the depth of detection and sensitivity of RGS has proven extremely valuable in identifying sentinel lymph node location, a lack of visual cues limits its ability to guide margin detection in real time.

FGS, on the contrary, provides real-time tumor visualization and excels in delineation of tumor borders and margins. Indocyanine green (ICG), a near-infrared (NIR) contrast agent, has seen recent expansion in its application, and has current utility in colorectal, urological, and oncological surgery. However, signal detection in FGS is unfortunately limited by overlying tissue, with minimal fluorescence detectable beyond a depth of 5 mm [10]. In surgical oncology, fluorescent visualization of tumors may also rely on the enhanced permeability and retention (EPR) effect, another nonspecific mechanism, through which increased ICG enters the tumor via abnormal, leaky vessels and is retained due to inadequate lymphatic drainage. The EPR effect remains, however, a nonspecific mechanism with inconsistent tumor uptake and potential for false positives, particularly with inflamed tissue, while still suffering from poor tissue signal penetration [11].

Due to the pitfalls of high rates of false positive and false negatives associated with passive accumulation, targeted fluorescent and radioactive agents have been designed to offer tumor and target specificity [7]. These newer, targeted agents exploit overexpressed tumor antigens or specific aspects of the tumor microenvironment. Pafolacianine (Cytalux), a folate receptor-targeting agent, has been FDA-approved for use in ovarian and lung cancer resection [10], and clinical trials are currently underway for FGS agents targeting the epidermal growth factor receptor (EGFR) in sarcoma and glioma resection, carcinoembryonic antigen (CEA) in colorectal cancer, and other epitopes. Pegulicianine (Lumisight) takes advantage of an acidic tumor microenvironment to provide specificity during lumpectomy for breast malignancies [12]. Similarly, targeted radiotracers are also being trialed in prostate cancer, neuroendocrine tumors, and other malignancies [13-15].

As such, combination of fluorescence, radioactivity, and targeted molecular modalities maximizes the strengths of each system. Indeed, development into dual-labeled targeted tracers has progressed with clinical trials of ^111^In-DTPA-labetuzumab-IRDye800 targeting CEA in colorectal cancer and ^111^In-DOTA-girentuximab-IRDye800 for carbonic anhydrase IX in renal cell carcinoma, and preclinical work evaluating ^111^In-DTPA-D2B-IRDye800CW for prostate surface membrane antigen in prostate cancer [16-19]. Given that none of these agents are intended for pediatric neuroblastoma and the need to improve surgical safety while performing an adequate resection, we sought to design and optimize a dual-labeled agent for clinical intraoperative use.

In neuroblastoma, GD2, a cell surface disialoganglioside, is overexpressed in 95-100% of tumors, but has only a limited presence in normal tissue [20], rendering it ideal for molecular targeting. Previous work from our lab demonstrated the feasibility of targeting GD2 for IMI in neuroblastoma using a nonclinical grade mouse-derived GD2 antibody [21]. To progress towards clinical translation, we sought to utilize Dinutuximab, an FDA-approved chimeric .antibody with the same variable region targeting GD2, as the antibody backbone for a clinically translatable IMI agent, and study its efficacy in more advanced clinically relevant rodent surgery. As such, to Dinutuximab, we conjugated IRDye800CW for near-infrared fluorescence detection and diethylenetriaminepentaacetic acid (DTPA) for chelation of Indium-111 for radiodetection, thus enabling dual-modality IMI. The near-infrared fluorophore IRDye800 has an excitation and emission profile similar to that of ICG [7], consequently enabling its detection across open, laparoscopic, and robotic surgery as near-infrared camera systems are already widely available. Its excitation and emission wavelengths in the near-infrared spectrum are also longer than that of visible light, thus minimizing overlapping background autofluorescence and absorbance from blood and hemoglobin [22]. Indium-111 is a gamma-emitting radioisotope with a half-life of 2.81 days, complementing the 1-4 week serum half-life of antibodies [23]. It is readily detected by commonly used intraoperative handheld devices, such as the Neoprobe^®^, and its energy emission profile is lower than that of other radioisotopes with similar half-lives [23], consequently allowing for lower overall radiation exposure to the patient and surgical staff. ^111^In-Dinutuximab-IRDye800 consequently has the potential to significantly improve pediatric neuroblastoma resection by enhancing real-time distinction between normal tissue and tumor, leading to a more complete resection and improved patient safety.

Here, we describe the development and optimization of ^111^In-Dinutuximab-IRDye800, including its chemical optimization, *in vitro* binding characterization, and *in vivo* performance mirroring clinical use.

## Methods

### Cell culture

SK-N-BE(2) (ATCC no. CRL-2271) cells were cultured in Dulbecco’s Modified Eagle Medium (Cat# BW12614F, Lonza) supplemented with 10% heat-inactivated FBS (Cat# SH3091003, Cytiva Hyclone™), 1x MEM NEAA (Cat# 11140-050, Gibco), 1x Antibiotic-Antimycotic (Cat #15-240-062, Gibco), 100 mM sodium pyruvate (Cat# BW13-115E, Lonza), 55 μM beta-mercaptoethanol (Cat# 21985023, Gibco), and 50 mg/mL Normicin (Cat# ant-nr-05, InvivoGen) and maintained in a 37°C humidified incubator with 5% CO_2_.

### Tracer synthesis

The final dual-labeled construct is ^111^In-DTPA-Dinutuximab-IRDye800 and will subsequently be referred throughout as ^111^In-Dinutuximab-IRDye800; further abbreviated forms are avoided to distinguish the antibody, chelator, radionuclide, and fluorophore components clearly. Dinutuximab in 0.9% sodium chloride solution (United Therapeutics) was adjusted to pH 8.5-9 using 0.1M sodium carbonate buffer. DTPA was conjugated by addition of a 25-fold molar excess of p-SCN-Bn-CHX-A’’-DTPA (Cat# B-351, Macrocyclics) and incubation at 37°C for 1 hour. Excess free DTPA was removed by centrifugal ultrafiltration using a 10 kDa molecular weight cutoff concentrator (Pierce Protein Concentrator, Thermo Fisher Scientific) with PBS as the diluent. IRDye800 conjugation was then performed by adding a 4-fold molar excess of IRDye800CW NHS ester (P/N 929-70021, LI-COR) at pH 8.5 – 9, followed by incubation at room temperature for approximately 30 minutes. DTPA and IRDye800 substitution levels were quantified by UV-Vis spectrophotometry (BioTek Epoch with Take3 plate, Agilent) using absorbance at 245 nm and 280 nm for DTPA, and 780 nm and 280 nm for IRDye800 (with a 3% correction for IRDye800 contribution at 280 nm). Conjugates were protected from light and stored at concentrations below 3 mg/mL until use.

DTPA-Dinutuximab-IRDye800 was radiolabeled with In-111 using a previously described method [24]. ¹¹¹In stock solution (BWXT) was diluted into 400 µL of sodium citrate buffer (pH 6) and allowed to equilibrate for 5 minutes prior to addition of 500-600 µg the conjugate. The mixture was incubated at 37°C for 1 hour, and radiolabeling efficiency was monitored by instant thin-layer chromatography (iTLC) using 50 mM sodium citrate for the mobile phase. The buffer was subsequently exchanged to DPBS (Cat# 175512F12, Lonza) by two washes with a total of 2 mL PBS using a Cytiva 30 kDa SEC filter membrane concentrator for administration via tail vein injection. Radiochemical purity was assessed by SEC-HPLC (BioSec3, Agilent; 3 µm, 300 Å, 4.6 × 300 mm column; PBS mobile phase, 0.4 mL/min), and purification was repeated until radiochemical purity exceeded 95%.

### ELISA

ELISA binding assays were performed to determine the impact of IRDye800 substitution level on the GD2-specific binding affinity of Dinutuximab. GD2 antigen was diluted in ethanol and added to wells of microtiter plates at 0.4 µg/well. After blocking with 1% BSA and rinsing, unmodified Dinutuximab or DTPA-Dinutuximab-IRDye800 (ratios of .9 DTPA and 1.55, 2.7, or 4.37 IRDye800 per antibody) were added to wells in triplicate at concentrations ranging from 2.6–333 nM, incubated, and rinsed. Next, the secondary antibody, HRP-conjugated goat anti-human IgG Fc (Thermo Fisher Scientific) was added, incubated, and rinsed, then HRP substrate was added. Wells containing immobilized GD2 and secondary antibody alone served as non-specific binding controls, and specific binding was obtained by subtraction from total binding values. Equilibrium dissociation constants (K_d_) and 95% confidence intervals were determined by non-linear regression using a one-site saturation binding model (GraphPad Prism 10).

### Fluorescence assessment

To characterize the impact of IRDye800 substitution level on fluorescent output, a dilution series with 5 concentrations of the above-mentioned conjugates (ratios of .9 DTPA and 1.55, 2.7, and 4.37 IRDye800 per antibody) was prepared within the previously established linear range of IRDye800 fluorescence. Fluorescence (excitation 760 nm, emission 789 nm) was measured using a plate reader (BioTek Epoch with Take3 plate; Agilent). The fluorescence intensity (FI) was plotted against antibody concentration for each conjugate, and linear regression was used to determine the fluorescence-per-antibody (FI/[Ab]) and fluorescence-per-IRDye800 (FI/[IRDye800] for each tracer.

### *In vivo* models of neuroblastoma

Animal studies were performed in accordance to the protocols approved by the University of Pittsburgh Institutional Animal Care and Use Committee (IACUC, protocol # 24013900). 4 to 6-week-old nude mice (Fox1nu, strain #002019; The Jackson Laboratory) and 4 to 6-week-old rats (RNU, strain 316, Charles River) were maintained in a temperature-controlled animal facility at the UPMC Children’s Hospital of Pittsburgh or the UPMC Hillman Cancer Center with a 12-hour light/dark cycle in cohorts of five mice per cage and two rats per cage. At least 48 hours passed between delivery of the animals and any procedures to allow for facility acclimation.

Orthotopic models of neuroblastoma were generated following previously established methods [21]. After anesthesia was achieved with inhaled isoflurane and confirmed via lack of pedal reflex, a left flank incision was made through skin, abdominal wall, and peritoneum to expose the left adrenal gland. After trypsinization, washing, and counting, 1 million or 2 million, for mice and rats respectively, human SK-N-BE(2) neuroblastoma cells were suspended in a mix of 20 µL of PBS and 20 µL of Matrigel (#354234; Corning), and directly injected into the left adrenal gland. The abdominal wall was closed with absorbable and the skin closed with surgical clips that were removed 10-14 days after surgery. Animals were placed in a clean cage on a warming pad and monitored until full recovery from anesthesia was noted. Accessible food and water were placed on and near the cage bottom. Tumors were grown for approximately four weeks and animals monitored at least weekly.

### Biodistribution studies

Rodents were used for biodistribution studies about four weeks after SK-N-BE(2) cells were injected, allowing for tumor growth to around 1 cm in size. For each experiment, mice or rats were randomly assigned to the described treatment groups. On the day of radiolabeling, ^111^In-Dinutuximab-IRDye800 was injected via tail vein (13.5-129.7 µCi/50 µg in 100 µL PBS for mice, 230 µCi/150 µg in 100 µL PBS for rats). Based on our prior work targeting GD2 for IMI in neuroblastoma [21], four days after tracer injection was selected as the timepoint for euthanasia and biodistribution analysis. Rodents were euthanized and organs (heart, lung, blood, right kidney, left kidney, liver, muscle, femur, small intestine, brain, contralateral adrenal, tumor, and spleen) were harvested and weighed. For gamma biodistribution, tissue-associated radioactivity was measured by gamma counter (PerkinElmer Wizard 2, model 2480; PerkinElmer) and quantified as percent injected dose per gram of tissue (%ID/g). This was adjusted for In-111 decay from time of tracer injection to time of measurement.

For optical biodistributions, the aforementioned tissues were placed on a minimally fluorescent black PLA printed plate, and images were captured with the SPY-PHI near-infrared camera (Stryker). These were saved using a NIX HDMI capture card (model USBC-CAP60, Plugable Technologies) and OBS video capture software (Open Broadcaster Software, open source). Images were windowed appropriately to allow for visualization of all organs and tissues. Regions of interest (ROIs) for each organ and the PLA plate background were drawn, then applied to the original unadulterated image (ImageJ, NIH). Mean fluorescence intensity (MFI) for each ROI was quantified in arbitrary units per squared pixel (AU/p^2^) and background fluorescence was subtracted.

### Dose determination

To determine the optimal number of IRDye800 molecules per antibody, DTPA-Dinutuximab-IRDye800 conjugates with differing IRDye800/Ab ratios (1.18, 1.72, 2.14, 2.71, and 3.09) were radiolabeled with Indium-111 then injected via the tail vein of neuroblastoma-bearing mice (n=4-6 per group). After 4 days, gamma and optical biodistributions were performed as described. After the optimal number of IRDye800 per antibody was determined, this process was repeated with varying doses of DTPA-Dinutuximab-IRDye800 based on concentration of Dinutuximab administered.

### Phototoxicity

300,000 SK-N-BE(2) neuroblastoma cells were incubated with 0, 5, 15, and 25 µg/mL of Dinutuximab-IRDye800 at a ratio of 1.4 IRDye800 per antibody for 5 hours. Treatment with hydrogen peroxide was used as a positive control for cell death. The Modulight system (ML8500, Modulight, Inc.) was then used to deliver broadband light at an irradiance of 100 mW/cm^2^, with doses of 3, 15, and 30 J/cm^2^ achieved by adjusting exposure duration (30 seconds, 150 seconds, and 300 seconds, respectively). Cell viability was assessed 24 hours post-irradiation by Annexin V/propidium iodide (PI) staining and flow cytometry.

### Clinically Significant Events study

To assess the intraoperative capabilities of ^111^In-Dinutuximab-IRDye800, the clinically significant events (CSE) study design was adapted from clinical trials of IMI agents and modified for the rodent setting. CSE were then defined as the incidence of tracer-guided identification of primary tumor not detected under white light, positive margins, or additional tumor deposits/synchronous lesions that were not previously identified during an attempt at full-resection under white light alone. Tumors were grown in 5-week-old RNU rats for 5 weeks (n=11), then DTPA-Dinutuximab-IRDye800 (150 µg in 150 µl PBS per rat) was administered via tail vein injection. After 4-5 days, surgical removal of tumors and any visible nodules was performed with standard-of-care white light, followed by IMI-guided surgery. The SPY-PHI near-infrared camera was used to capture images prior to any resection, then intraoperatively to identify and resect any residual fluorescent tissue, regardless of clinical correlation to malignant-appearing tissue. Rats were euthanized and the residual tumor bed was also resected. All resected tissue (primary tumor, nodules, fluorescent tissue, and residual tumor bed) was sent for blinded histopathologic analysis. The total number of CSE and percent of rats with CSE were calculated, along with the sensitivity, specificity, positive predictive value (PPV), and negative predictive value (NPV).

### Statistical analysis

Graphs were generated and statistical analysis were performed using GraphPad Prism v11.0.2. Comparisons across multiple groups were performed using one-way or two-way ANOVA, as appropriate, with post-hoc multiple comparison testing where indicated. Data is presented as mean ± SEM unless otherwise specified. A two-sided p-value < 0.05 was considered statistically significant. Animals that died immediately after injection because of formulation intolerance were excluded from biodistribution analysis but were reported as tolerability outcomes.

## Results

### Synthesis and optimization of ^111^In-Dinutuximab-IRDye800

#### Binding

Unmodified Dinutuximab bound to GD2 with a dissociation constant (K_d_) of 6.89 nM (95% CI: 5.40–8.76 nM). DTPA-Dinutuximab-IR800 was synthesized with approximately 1 DTPA and a range of 1.55-4.37 IRDye800 per antibody. Binding affinity was largely preserved with 0.9 DTPA and 1.55 IRDye800 per antibody (K_d_ 5.24 nM, 95% CI: 4.39–6.26 nM) and slightly reduced with 0.9 DTPA and 2.70 IRDye800 per antibody (K_d_ 18.63 nM, 95% CI: 15.08–22.95 nM). However, maintaining 0.9 DTPA per antibody but increasing the substitution level to 4.37 IRDye800 per antibody resulted in a substantial loss of binding affinity, with K_d_ 75.79 nM (95% CI: 54.16–105.3 nM, Figure 1a).

**Figure 1.**
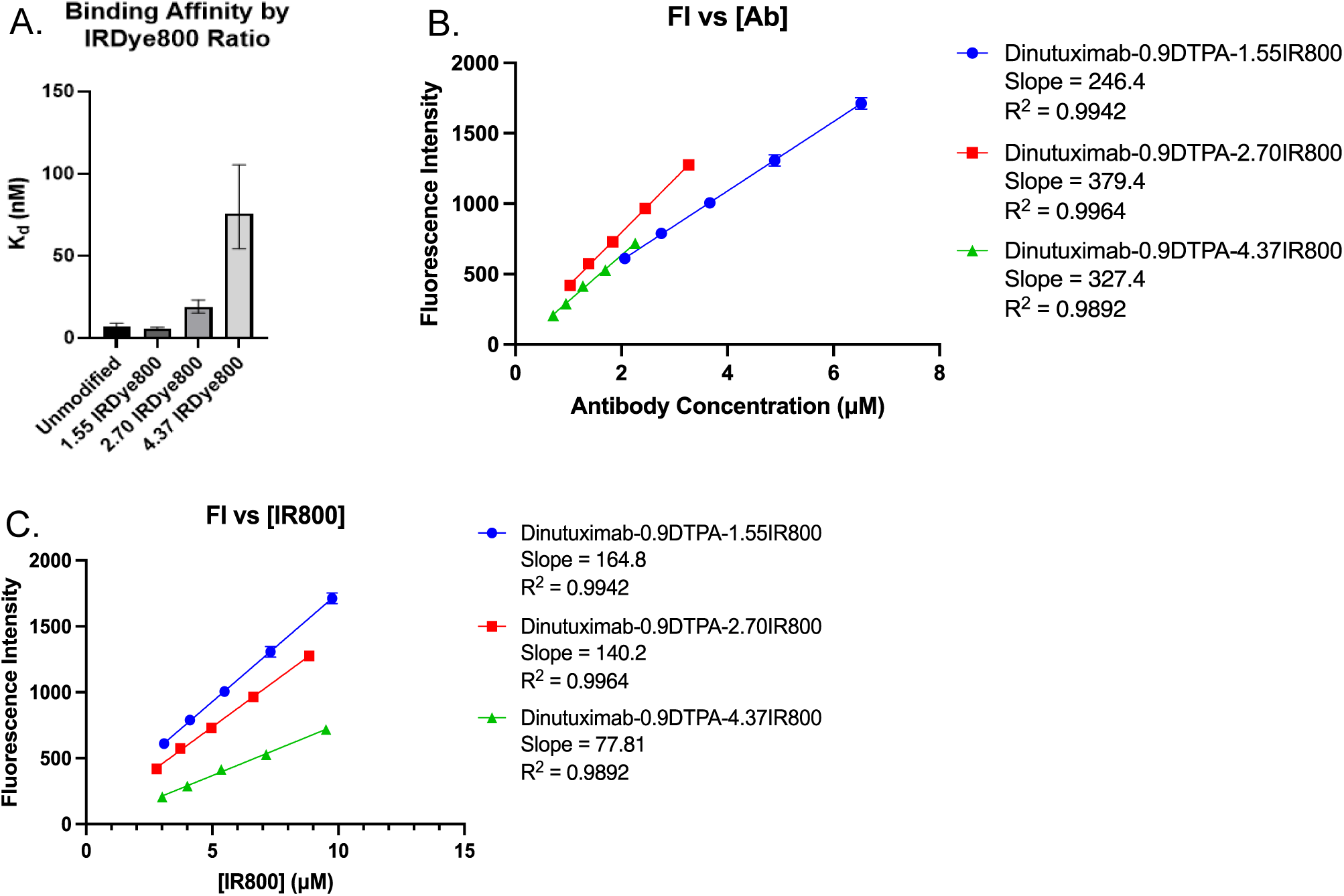
(a) Binding affinity (K_d_) of Dinutuximab-based tracer decreases with increasing IRDye800 ratios; error bars represent 95% confidence intervals. (b) Fluorescent output of IRDye800 peaks at 2.70 IRDye800 per antibody. When plotted against antibody concentration (b) and IRDye800 molar concentration (c), fluorescence intensity does not demonstrate the same increase between 2.70 and 4.37 IRDye800/antibody as it did between 1.55 and 2.70 IRDye800 per antibody. Self-quenching was observed at higher dye-to-antibody ratios.

#### Fluorescence intensity

Fluorescence intensity per tracer initially increased, from 246.4 AU/µM with 1.55 IRDye800 per tracer to 379.4 AU/µM with 2.70 IRDye800 per tracer, a 1.54-fold increase. Additional fluorophore did not increase fluorescence, remaining at 327.4 AU/µM with 4.37 IRDye800 per antibody, suggesting a component of quenching at higher ratios (Figure 1b). Linear regression of fluorescence intensity versus antibody concentration had an excellent fit (R^2^ = 0.9942, 0.9964, and 0.9892 for 1.55, 2.70, and 4.37 IRDye800/antibody, respectively). Similarly, fluorescence per IRDye800 was highest at the lowest substitution level (164.8 AU/µM at 1.55 IRDye800 per antibody) and declined with increasing substitution levels (140.2 and 77.81 AU/µM at 2.70 and 4.37 IRDye800 per antibody, respectively; Figure 1c), consistent with self-quenching of IRDye800 at higher dye-to-antibody ratios.

#### Impact of IRDye800 substitution level on biodistribution

In determining the optimal IRDye800 degree of labeling, neuroblastoma-bearing mice were administered ^111^In-Dinutuximab-IRDye800 with 1.32 DTPA and a range of IRDye800 per antibody (1.18, 1.72, 2.14, 2.71, and 3.09). Higher degrees of labeling appeared to increase the risk of formulation instability. In the 3.09 IRDye800 group, all mice were found to have died shortly after injection despite adequate recovery from tracer injection and anesthesia. Tracer injections were subsequently slowed and each mouse monitored for full recovery from anesthesia prior to advancing to the next mouse. In the 2.71 IRDye800 group, 1 mouse died after injection and 2 required cardiopulmonary resuscitation, but recovered fully. For both groups, a visible ring of particulate material was noted afterwards in the injection tubing, suggesting aggregation of the compound that may have led to the observed toxicity. Substitution levels of 1.18, 1.72, 2.14 IRDye800 per antibody were well-tolerated without any observed adverse reactions or aggregation.

Tumor selectivity of the tracer was statistically unaffected by degree of optical labeling (7.0-10.8%ID/g for 1.18-2.71 F/Ab; p=ns). However, with increased IRDye800 per antibody ratios, progressive increases in off-target hepatic accumulation of the tracer were observed from both optical (8.6, 10.1, 13.2, and 14.3 AU/p^2^, respectively; p<0.0001; Figure 2a) and gamma standpoints (2.09, 5.55, 7.42, and 8.81%ID/g; p<0.0001; Figure 2b). A ratio of 1.18 IRDye800 per antibody ultimately demonstrated the optimal combination of gross fluorescent signal and tumor-to-background ratio with respect to the muscle (14.8), blood (13.6), and liver (4.5).

**Figure 2.**
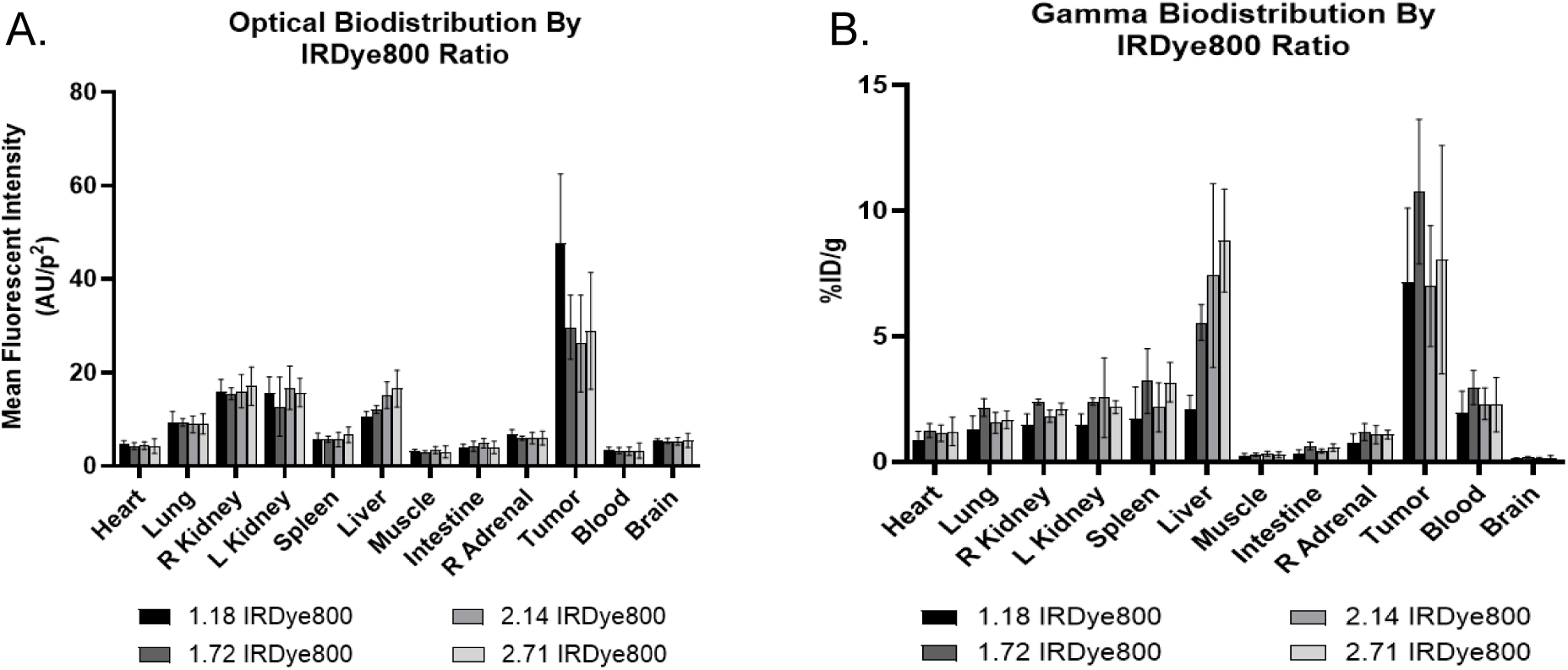
IRDye800 ratios impact off-target accumulation of ^111^In-Dinutuximab-IRDye800 and inform the optimal fluorescent imaging window. (a) Tumor-specific uptake of ^111^In-Dinutuximab-IRDye800 by gamma biodistribution was comparable across all ratios. (b) Optimal tumor-to-background ratio was obtained with 1.18 IRDye800 per antibody. Increased off-target hepatic accumulation was observed with increased IRDye800 per antibody from both a gamma and fluorescent perspective.

#### Optimization of dose and timing

Neuroblastoma-bearing mice were injected with 5 µg (5 µCi), 15 µg (20 µCi), and 45 µg (30 µCi) of ^111^In-Dinutuximab-IRDye800 in 100 µL DPBS (n=5 per group). All injected doses were tolerated without sign of any adverse effects. Optical and gamma biodistribution studies performed 4 days after tracer injection demonstrated specific accumulation of the tracer in the tumor. Optical biodistribution demonstrated a dose-dependent increase of NIR visibility and detection, reflecting a potential detection threshold of the SPY-PHI camera for IRDye800 (Figure 3a). No significant differences between tumor (3.88±0.74 AU/p^2^) and other resected organs were detected with a 5 µg dose. With 15 ug, while tumor uptake (7.04±1.43 AU/p^2^) was significantly higher than all other resected organs (p<0.01), these differences were not easily visible on raw NIR images. However, a higher dose of 45 µg of tracer demonstrated both significantly higher tumor uptake (24.9±8.28 AU/p^2^, p<0.01) on NIR image analysis as well as raw images from the handheld SPY-PHI camera. Gamma biodistribution demonstrated that regardless of dose, relative accumulation of the tracer in tumor remained similar (35.0±17.3%ID/g, 27.4±6.58%ID/g, and 32.0±11.1%ID/g for 5 µg, 15 µg, and 45 µg, respectively, Figure 3b). Furthermore, tumor and off-target tracer accumulation was not impacted by higher degrees of ^111^In activity. Activities of 25, 90, and 130 µCi per 50 µg ^111^In-Dinutuximab-IRDye800 demonstrated similar tumor uptake in terms of %ID/g (19.4%, 21.8%, and 20.8%, respectively, Figure 3c).

**Figure 3.**
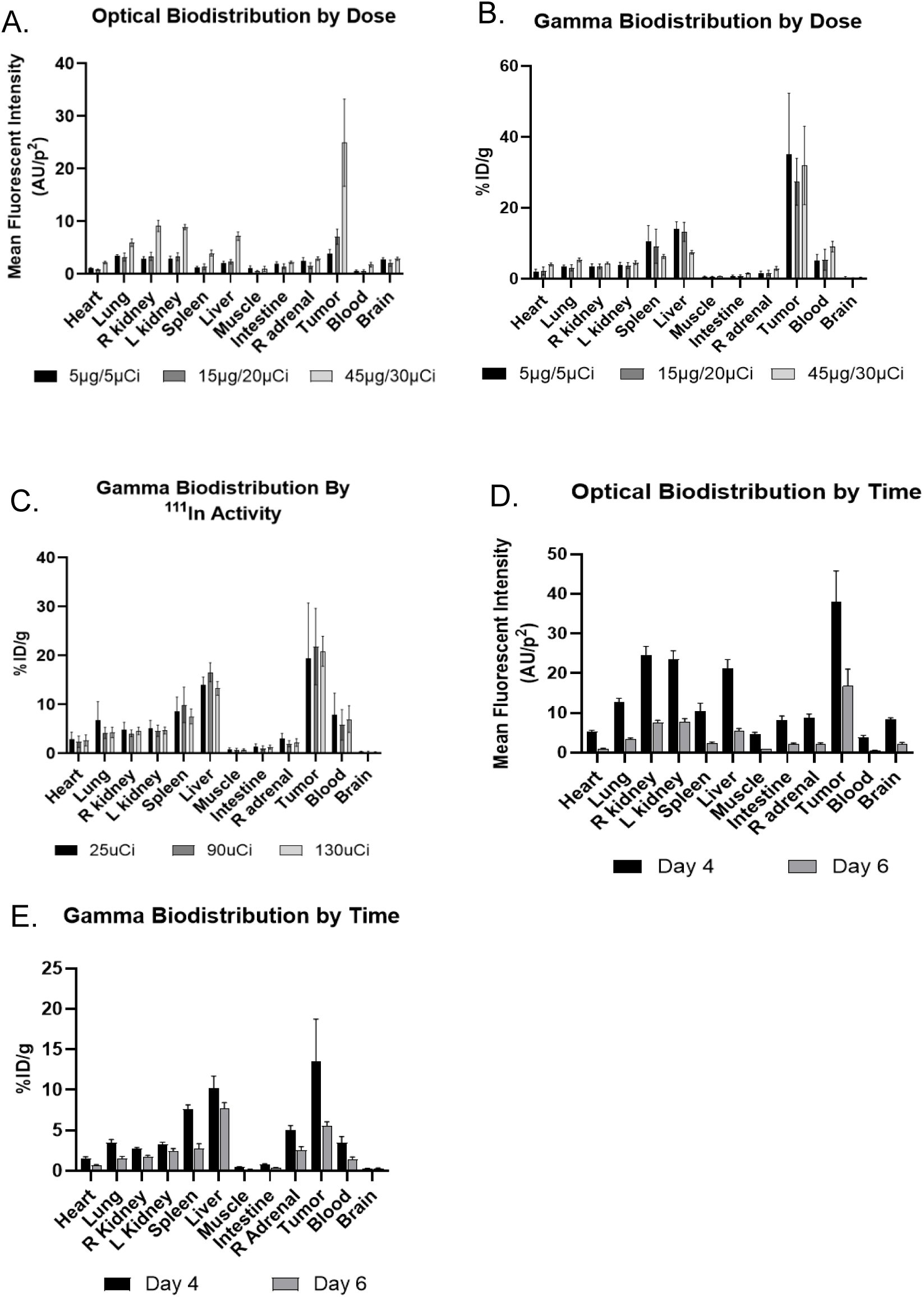
Optimization of dose and timing demonstrates that 45-50 µg of tracer assessed at 4 days after tracer administration are the optimal parameters for use of ^111^In-Dinutuximab-IRDye800. (a) Optical biodistribution demonstrates a dose-dependent NIR signal where 45 µg of tracer produced both visible fluorescence on raw SPY-PHI images and statistically significant tumor uptake of ^111^In-Dinutuximab-IRDye800. (b) Gamma biodistribution across the same three doses demonstrating comparable tumor uptake of the tracer as measured by %ID/g. (c) Differing levels of ^111^In activity of ^111^In-Dinutuximab-IRDye800 does not impact its specificity for neuroblastoma tumor. Optical (d) and gamma (e) biodistribution demonstrate that 4 days after tracer administration is the optimal period for fluorescent and radioactive assessment given the observed global decreases in MFI and %ID/g on Day 6.

Both 4 and 6 days after tracer injection were evaluated to determine the ideal time frame for future surgery. 50 µg of ^111^In-Dinutuximab-IRDye800 was administered, and fluorescent and gamma biodistributions performed at each timepoint. Specificity of ^111^In-Dinutuximab-IRDye800 was again demonstrated at both timepoints. Average fluorescence intensity of the tumor was significantly higher (37.9 AU/p^2^ on Day 4 and 16.8 AU/p^2^ on Day 6) than all other measured organs (p<0.0001). However, Day 4 yielded a higher mean fluorescence intensity in the tumor than on Day 6 (37.9 AU/p^2^ vs 16.8 AU/P^2^, p<0.0001). There was an overall decreased fluorescence intensity observed in all organs on Day 6 compared to Day 4 (Figure 3d). On gamma biodistribution, the impact of time was noticeably more pronounced (Figure 3e). Between days, %ID/g of the tumor fell from 13.5% to 5.58% (p=0.0002). Additionally, on Day 4, tumor uptake of ^111^In-Dinutuximab-IRDye800 (13.5%ID/g) was significantly higher than all other measured organs (p<0.02) aside from liver (10.2% ID/g, p=0.69) and spleen (7.66% ID/g, p=0.09). Tumor-to-blood ratio was 3.86 (blood 3.49%ID/g) and tumor-to-muscle ratio was 30.2 (muscle 0.45%ID/g). On Day 6, tumor uptake of ^111^In-Dinutuximab-IRDye800 (5.58 %ID/g) was significantly lower than that of the liver (7.70 %ID/g, p<0.001). Tumor-to-blood ratio was similar to Day 4 at 3.88 (blood 1.44 %ID/g) and tumor-to-muscle ratio 29.1 (muscle 0.19%ID/g). Similar to fluorescence intensity, although tumor %ID/g was higher than other measured organs except the liver, there was globally decreased radioactive signal in all measured tissues.

#### Phototoxicity assessment

Given that IRDye800 is a core component of this Dinutuximab -based tracer, we sought to evaluate the potential for phototoxic effects from the presence of the near-infrared fluorophore itself. Flow cytometry with Annexin V and propidium iodide as markers of cell death and apoptosis demonstrated largely unchanged cell viability across all tracer concentrations and light doses, with percent remaining of viable cells ranged from 70.4% to 85.9%, compared to 74.2% in the untreated control (Figure 4). There were no significant differences in cell viability due to tracer dose, light dose, or any combination thereof compared to no treatment, including cells exposed to the highest tracer concentration of 25 µg/mL (76.3% viability, p=ns), a supraclinical light dose of 30 J/cm² (85.9% viability, p=ns), and a combination of both (71.8% viability, p=ns, Figure 4). In contrast, treatment with hydrogen peroxide, used as a positive control for cell death, demonstrated 16% cell viability.

**Figure 4.**
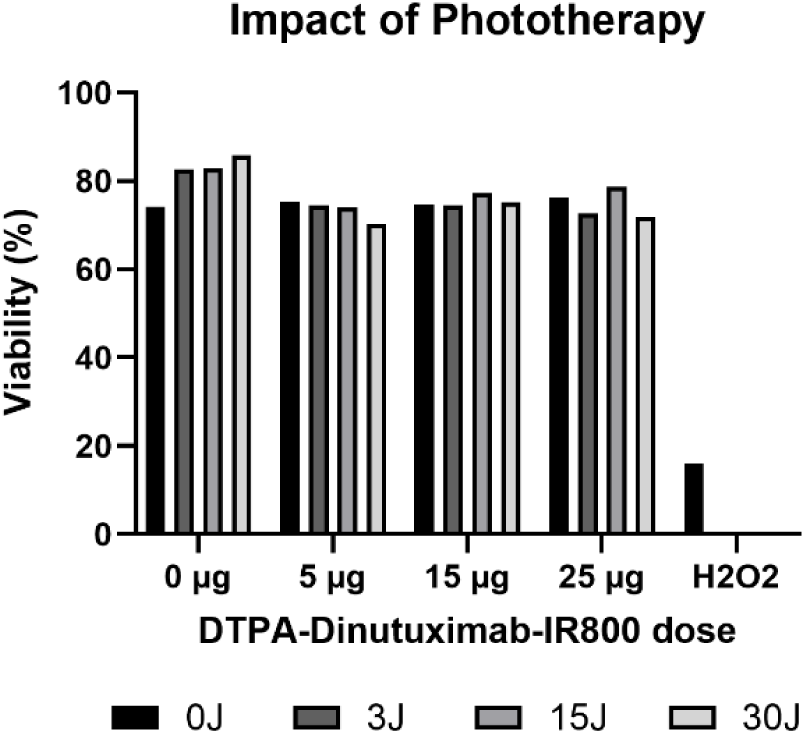
DTPA-Dinutuximab-IRDye800 does not induce phototoxicity at or beyond clinically relevant concentrations (in µg/mL) of Dinutuximab or light doses. Cell viability did not significantly differ across all treatment conditions in comparison to the negative control of 0 µg of tracer and 0 J of light treatment.

### Tracer Specificity

After determination of dose, degree of optical labeling, and time interval after injection, a biodistribution of ^111^In-Dinutuximab-IRDye800 was performed and compared to isotype control tracer (^111^In-isotype-IRDye800) in both male and female mice. ^111^In-Dinutuximab-IRDye800 demonstrated substantially higher tumor specificity and lower nonspecific accumulation, observed in both optical and gamma biodistributions. This was also observed across male and female mice without significant differences between the two groups. Fluorescence biodistribution analysis (Figure 5a) with 2-way ANOVA demonstrated a significantly higher mean fluorescence intensity of the tumor (male: 24.42 AU/p^2^, female: 25.45 AU/p^2^) in mice receiving ^111^In-Dinutuximab-IRDye800 compared to the isotype control (male: 10.92 AU/p^2^, female: 5.80 AU/p^2^, p<0.0001). Similar to optimization experiments, tumor fluorescence with ^111^In-Dinutuximab-IRDye800 on gross images (Figure 6) and fluorescent analysis was significantly higher than all other tissues in both groups (p<0.0001). Mean fluorescent intensities of particular interest were the ipsilateral left kidney (male: 2.57 AU/p^2^, female: 6.80 AU/p^2^); liver (male: 3.77 AU/p^2^, female: 4.46 AU/p^2^); muscle (male: 0.60 AU/p^2^, female: 0.89 AU/p^2^); and blood (male: 0.40 AU/p^2^, female: 0.73 AU/p^2^). Averaged between both sexes, tumor-to-kidney ratio was 6.62, tumor-to-muscle ratio was 34.65, and tumor-to-blood ratio was 47.96.

**Figure 5.**
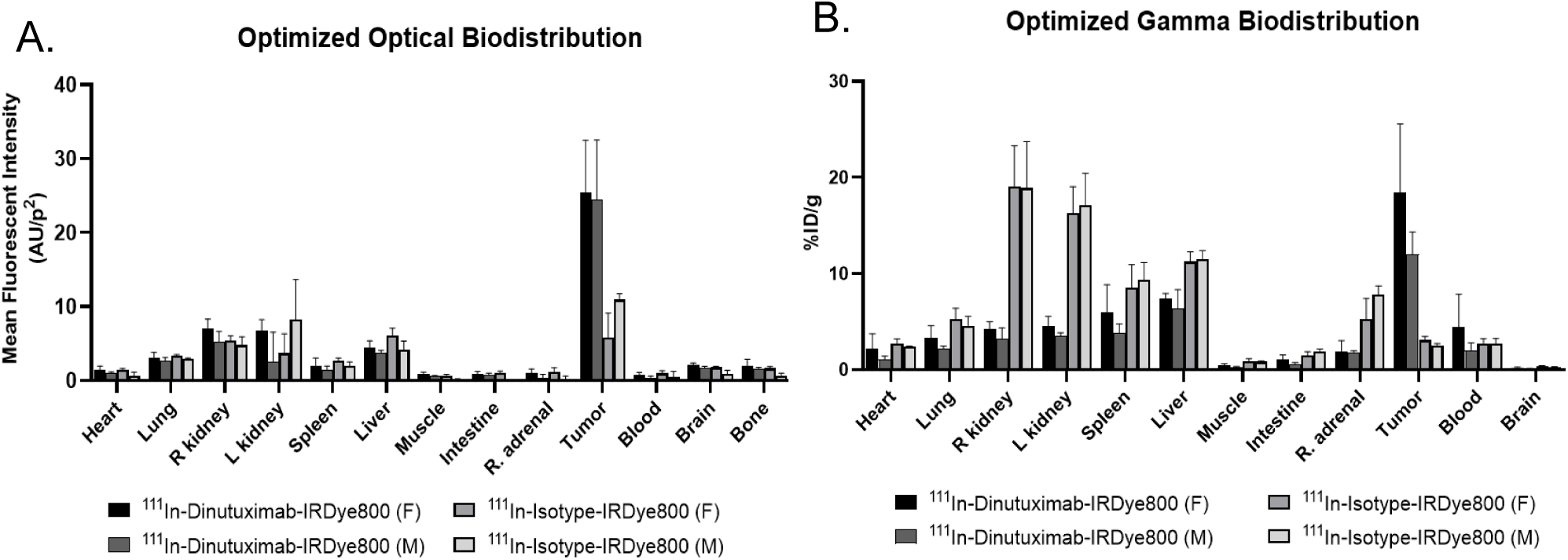
Optimized ^111^In-Dinutuximab-IRDye800 demonstrates tumor-specific accumulation compared to isotype control. (a) Mean fluorescence intensity of the tumor is significantly higher than all other resected organs and tissue (p<0.0001). (b). %ID/g of the tumor is significantly higher than all other organs and tissue (p<0.0001) whereas accumulation of the isotype control tracer was lower in the tumor than other organs such as kidney, spleen, and liver.

**Figure 6.**
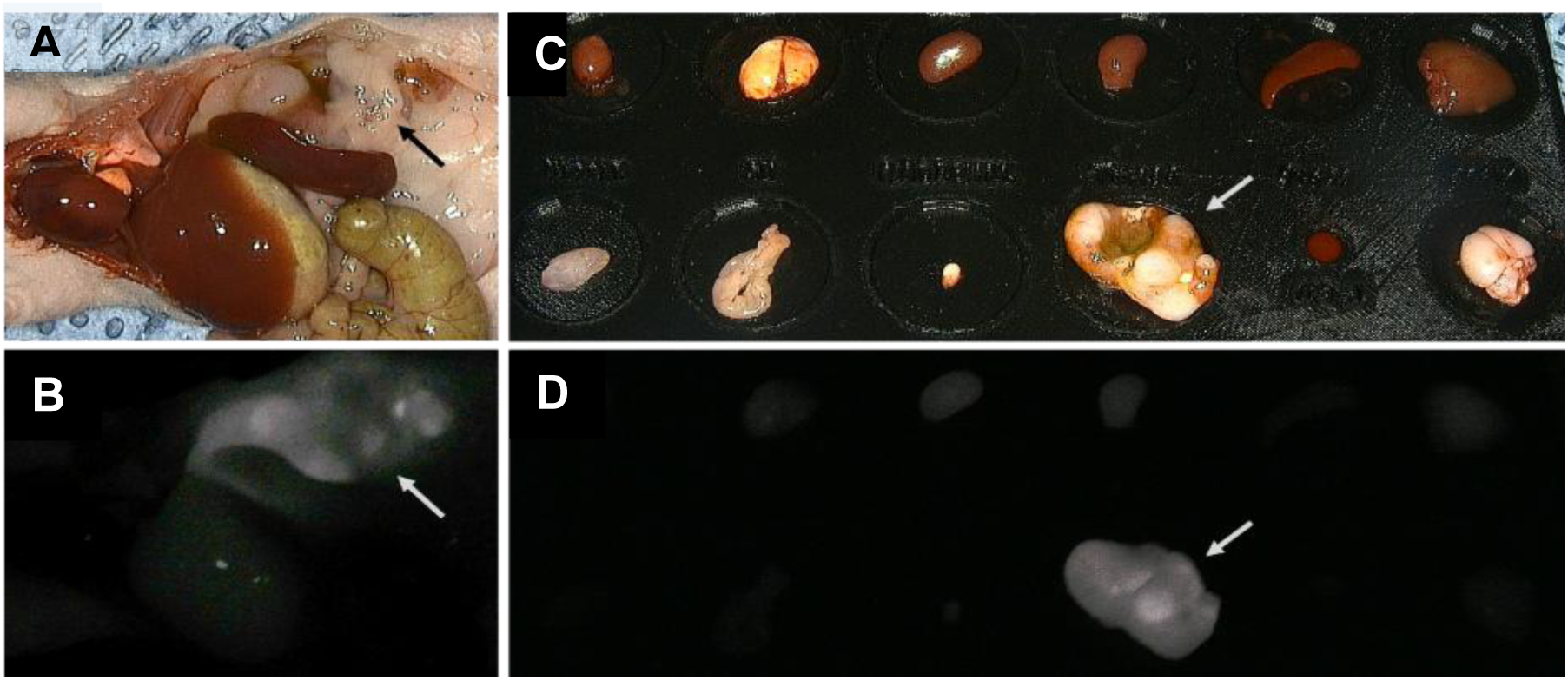
^111^In-Dinutuximab-IRDye800 yields tumor-specific fluorescence in comparison to other resected organs and tissues. Representative images of the tumor (arrow) *in situ* under white light (a) and the corresponding fluorescent image under NIR light (b), demonstrating strong tumor signal against background tissues. (c) White-light image of resected tumor and organs on a PLA plate for *ex vivo* biodistribution analysis with the corresponding NIR fluorescent image (d), again redemonstrating strong tumor-specific signal.

Similarly, %ID/g of the tumor was significantly higher in mice receiving ^111^In-Dinutuximab-IRDye800 (male: 12.0, female: 18.44), compared to ^111^In-isotype-IRDye800 (male: 2.54, female: 3.07, p<0.0001). In both male and female mice who received ^111^In-Dinutuximab-IRDye800, %ID/g of the tumor was also significantly high than all other organs (Figure 5b, p<0.0001). This was notable compared to muscle (male: 0.25%ID/g, female: .44%ID/g), blood (male: 2.03%ID/g, female: 4.34%ID/g), and ipsilateral left kidney (male: 3.53%ID/g, female: 4.50%ID/g). Combining results from both sexes, tumor-to-muscle ratio was 44.95, tumor-to-blood 5.05, and tumor-to-kidney 3.75.

### A study of clinically significant events

To assess the *in vivo* performance of ^111^In-Dinutuximab-IRDye800, a clinically significant events study design was modeled after clinical trials of IMI agents and modified for the rodent setting. Given their larger size, a rat model, as previously described [25], was utilized instead of a mouse model. Eleven neuroblastoma-bearing rats were injected with DTPA-Dinutuximab-IRDye800 (1.08 DTPA and 0.99 IRDye800 per antibody per tracer). All tumors, suspicious nodules, residual fluorescent tissue, and tumor beds were resected 4-5 days after injection. A total of 11 primary tumors and 16 additional nodules were resected under standard-of-care white-light surgery (Figure 7). These were all confirmed on pathology to be neuroblastoma. Of these, all primary tumors (100%) and 13 of the 16 nodules (81%) were also fluorescent. Fluorescence-guided surgery was then performed for each rat, leading to the resection of 20 additional nodules, 14 of which (60%) were confirmed neuroblastoma on pathologic analysis. FGS was able to detect lesions as small as 1.5 mm, with 4 of the additionally resected nodules under 2 mm in diameter. In 7 of the 11 rats (64%), fluorescence identified at least one additional malignant lesion that was not seen with normal white light. 9 of the 11 rats (82%) had no evidence of residual disease in the tumor bed after fluorescence-guided resection. Collectively, fluorescence-guided surgery with DTPA-Dinutuximab-IRDye800 for neuroblastoma had a sensitivity of 88%, specificity of 60%, PPV of 86%, and NPV of 64%.

**Figure 7.**
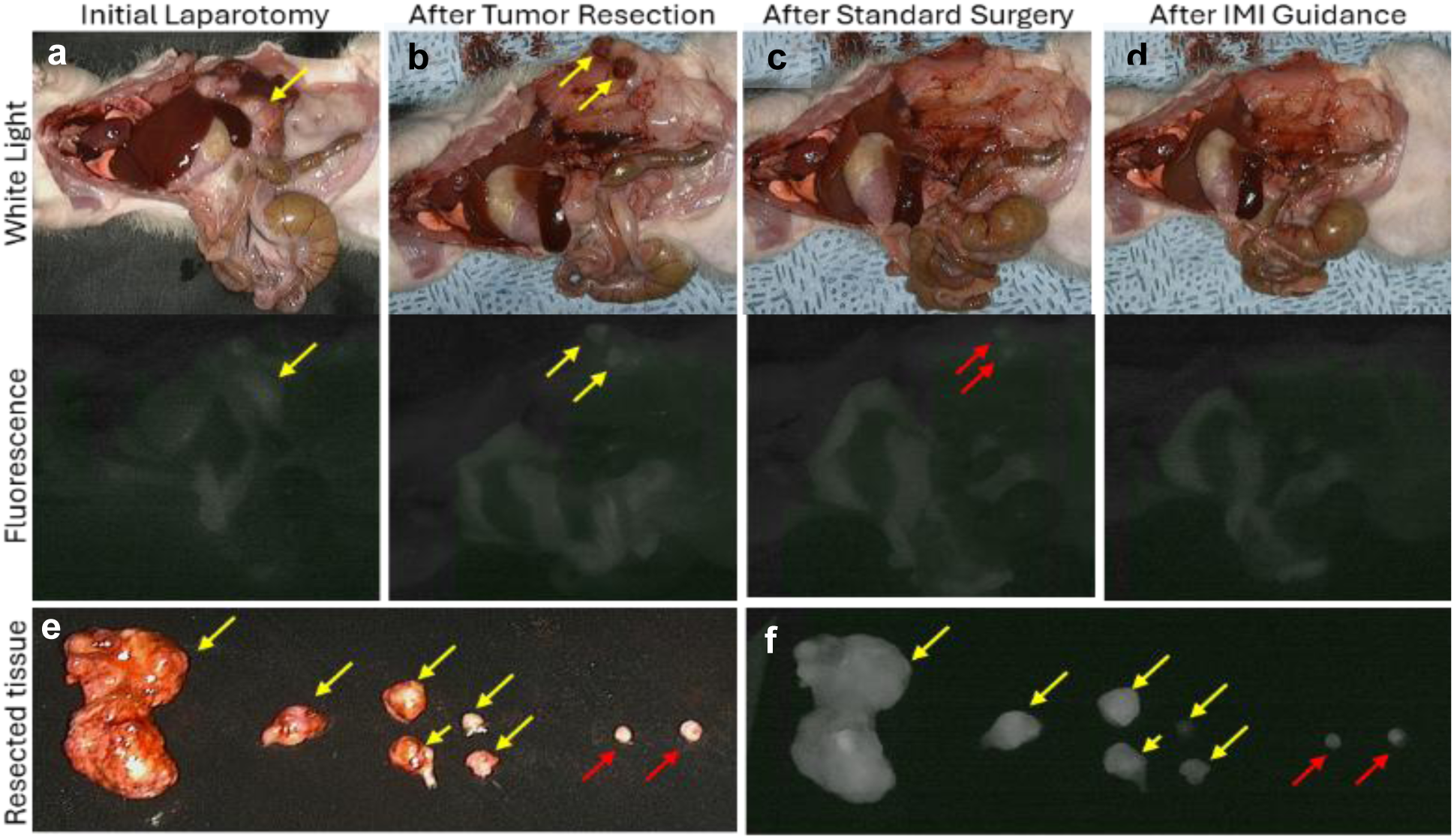
Fluorescence-guided surgery with ^111^In-Dinutuximab-IRDye800 can detect malignant disease not identified under normal white light. Representative images of standard-of-care white light (top row) and fluorescence-guided (middle row) surgical resections of neuroblastoma in a rat model. (A) The primary tumor was first resected without the use of FGS. (B) Additional suspicious nodules were identified without FGS and resected. (C) Additional nodules were then identified with FGS and resected. (D) Once all visible and fluorescent tumor was resected, the tumor bed was also resected and all specimens were sent for pathology. (E) Primary tumor and white light-visible nodules (yellow arrows), as well as fluorescence-visualized nodules (red arrows) were (F) fluorescent.

## Discussion

Here, we present the development and optimization of ^111^In-Dinutuximab-IRDye800, a novel dual-modality IMI agent intended for use in pediatric neuroblastoma resection. A central challenge in the design of antibody-based imaging conjugates is preserving the targeting affinity of the parent antibody while achieving sufficient signal for intraoperative detection. With the addition of multiple elements to the parent antibody, there may be physical as well as chemical changes that impact its overall binding affinity and function. Through systematic characterization of binding affinity, fluorescence intensity, and *in vivo* biodistribution, we optimized the tracer for tumor-selective accumulation, fluorescent and gamma signal, and meaningful intraoperative guidance in rodent models of neuroblastoma. Together, these findings support the translation of ^111^In-Dinutuximab-IRDye800 into the clinical setting.

### IRDye800 Optimization

While up to 1 DTPA and 1.6 IRDye800 per antibody did not significantly impact the binding affinity of the tracer, higher IRDye800 substitution levels (2.7 and 4.37 IRDye800 per antibody) were noted to lead to decreased binding affinity. Additionally, there was limited observed return on fluorescence intensity, with the implication that simply increasing dye per antibody ratios does not yield more useful or brighter signals. The conjugation of large molecules to antibodies is known to result in reduced binding to surface antigens, likely due to steric hindrance [26]. This was observed in our results as the highest degree of optical labeling of 4.37 IRDye800/antibody yielded a nearly 4-fold lower binding affinity than the next highest level of 2.7, and 15-fold lower affinity than 1.55, the degree of labeling that was closest to the physiologic binding affinity of Dinutuximab alone.

Compounding the loss of binding affinity, the phenomenon of self-quenching is an established complication of fluorescent dye use, in which a reduction in emission intensity occurs as a result of interactions between fluorescent molecules themselves [27]. Our results demonstrated that fluorescence intensity per tracer did not increase with additional IRDye800 molecules beyond a ratio of 2.7 per antibody. As the molecular structure of IRDye800 as a highly lipophilic and polarizable molecule, higher degrees of labeling can increase the number of coupled fluorophores and increase the amount of nonradiative molecular relaxation, ultimately reducing fluorescent emission [28]. This was also observed with our assessment of IRDye800 substitution levels as per-antibody fluorescence intensity progressively decreased as substitution level increased. Higher degrees of labeling may also increase the potential for dye-mediated antibody aggregation, which could explain the increased toxicity observed during *in vivo* injections of ^111^In-Dinutuximab-IRDye800 with 3.09 and 2.73 dyes/antibody. From the chelator DTPA standpoint, higher substitution levels were also noted to have detrimental effect on the functionality of ^111^In-Dinutuximab-IRDye800. Altogether, the impact of conjugation on binding affinity, fluorescence intensity, and tracer aggregation demonstrated a dye-to-antibody ratio approximately 1-2 IRDye800/antibody provided effective and adequate signal without compromised function or observed toxicity to cells.

### Dosage and timing

The next important considerations in the optimization of ^111^In-Dinutuximab-IRDye800 were the dosage and duration between tracer administration and radiofluorescent evaluation. Our dose-finding data highlighted that while lower doses of 5 µg and 15 µg demonstrated preferential tumor uptake on gamma and fluorescent biodistribution analysis, these doses were too low to yield sufficient fluorescent signal for a commonly used intraoperative NIR camera, limiting the benefit of one arm of our dual-modality tracer. Optical biodistribution analysis was performed through post-hoc windowing of still images captured by the near-infrared camera, analogous to adjusting the tissue windows on CT scans after the initial image is obtained. This underscores that while the selectivity of ^111^In-Dinutuximab-IRDye800 for neuroblastoma is maintained at low doses, detectable real-time signal must be the standard for intraoperative guidance. A dose of 45 µg produced both statistical and real-time visually detectable signal, and 50 µg was selected as the standard dose for subsequent experiments to provide a conservative margin above this threshold.

For an IMI agent to successfully distinguish between its target tissue of interest and normal tissue, it must be cleared from blood and non-target tissues such that tumor-to-background contrast is high enough for reliable detection [26]. By 4 days after tracer administration, mean fluorescence intensity and %ID/g of the tumor were statistically and grossly higher than that of blood and other surrounding tissue. At Day 4 and Day 6, there was a statistically significant increase in MFI of the tumor compared to all other organs and tissues. However, Day 6 had overall decreased MFI in all tissues despite tumor-to-blood and tumor-to-muscle ratios that appeared improved from Day 4 (39.1 vs 9.98, 20.7 vs 8.29, respectively). Following a similar trend, on Day 4, tumor uptake of the tracer by gamma biodistribution was significantly higher in all tissues other than liver and spleen. However, by Day 6, overall signal was decreased with markedly greater signal in the liver than the tumor itself. From a radioactivity standpoint, tumor-to-blood (3.86 Day 4 vs. 3.88 Day 6) and tumor-to-muscle (30.2 Day 4 vs 29.1 Day 6) ratios remained comparable. However, the globally decreased radiofluorescent signal of Day 6 indicate that metabolism continued to the point where absolute tumor signal was substantially diminished. Taken altogether, a dose of 50 µg with an optical degree of labeling between 1-2 IRDye800 per antibody, chelator ratio of 1-1.5 DTPA per antibody, and assessed at 4 days after administration provided the strongest and most tumor-specific signal.

In positioning ^111^In-Dinutuximab-IRDye800 for clinical application, comparison to an isotype-matched control confirmed specific GD2-driven tumor targeting of ^111^In-Dinutuximab-IRDye800. Upon combination of all optimized factors related to the tracer itself, ^111^In-Dinutuximab-IRDye800 yielded favorable tumor-to-background ratios, particularly against blood, muscle, kidney, and intestines, all of which may be encountered during neuroblastoma resection, helping to distinguish tumor from surrounding tissues and potentially identify occult deposits of disease. Although high hepatic and splenic background accumulation was noted, this is to be expected with IgG-based compounds [29], and poses less of an interpretative challenge during surgery as the tumors are typically not involving those organs.

In applying ^111^In-Dinutuximab-IRDye800 *in vivo,* the clinically significant events study (CSE) was designed to mirror the structure of clinical IMI trials and provide a more surgically relevant assessment of ^111^In-Dinutuximab-IRDye800. As such, CSE were defined as the incidence of tracer-guided identification of primary tumor, positive margins, or additional tumor deposits/synchronous lesions that were not previously identified during an attempt at full-resection under white light alone [30, 31]. Additional malignant lesions were detected with IMI guidance in 64% of animals. Compared to the CSE rates of pafolacianine in ovarian and lung cancer, 33% and 53% respectively, this suggests that ^111^In-Dinutuximab-IRDye800 also has the potential to positively influence intraoperative decision making. Detection of lesions as small as 1.5 mm also becomes clinically relevant as these subcentimeter deposits of disease may not be reliably identified on preoperative imaging or through manual palpation. Together, these findings suggest that the tracer can identify surgically occult disease deposits in a manner analogous to clinically significant events reported in human IMI studies.

The specificity of 60% warrants direct consideration as it does not completely reflect standard intraoperative decision making. As this study was not a survival study, any tracer positive tissue was resected regardless of gross surgical suspicion, which likely impacted the observed false-positive rate, and artificially decreased tracer specificity. While ^111^In-Dinutuximab-IRDye800, as all imaging tracers, is subject to some inherent false-positive signal, any intraoperative radiofluorescent signal would have been interpreted along with the context of the positive tissue itself. In practice, the surgeon would integrate positive signals with manual palpation, preoperative imaging, and clinical experience that would altogether serve to enhance effective intraoperative specificity. After any additional tissue was resected under IMI guidance, the tumor bed was also resected and sent to pathology to look for “missed” residual disease. This also does not mirror true surgical practice as these rats were euthanized after all samples were obtained, allowing for more aggressive and extensive tissue resection. However, this facilitated a more thorough characterization of the intraoperative capabilities of ^111^In-Dinutuximab-IRDye800.

The dual-modality design of ^111^In-Dinutuximab-IRDye800 is intended to combine the complementary systems of visual guidance and depth information, overcoming the inability of near-infrared fluorescence to penetrate more than 5 mm of tissue. Our group previously described the limits of detection of our dual-labeled tracer, using standard intraoperative tools to detect lesions buried within porcine organs and tissue phantoms [32]. We demonstrated superior detection of gamma signal compared with fluorescence in lesions deeper than 4-5 mm. In fact, gamma signal could be reliably detected in those models as deep as 4 cm. However, in a rodent model, there is very limited depth to any tissue, organ, or structure, thus limiting the conditions under which the depth advantage of radioactivity can be meaningfully assessed. As such, we decided to focus on fluorescence guidance in the CSE study, as we have previously described the clear benefit of radioguidance in detecting deeper tracer signal using porcine organs and tissue phantoms [32]. Our prior rodent work has also demonstrated that gamma decay signal, as detected by the handheld Neoprobe, was significantly higher in the tumor-bearing left flank of mice and rats compared to all other measured body regions, including chest, tail, and hind limbs [21]. Given that the radioactive component of the tracer supports deeper lesion detection and the limited size of the rodent abdominal cavity, future large-animal or clinically scaled studies will be useful to further define the degree to which intraoperative gamma detection complements fluorescence in anatomically complex resections.

## Limitations

We recognize several limitations in this study. All experiments in this study were conducted using a single human neuroblastoma cell line to generate orthotopic xenograft models in immunocompromised rodents. Consequently, this may not fully capture the tumor heterogeneity that exists in pediatric neuroblastoma or the pharmacokinetic implications of an intact immune system. However, GD2 expression, while variable, is nearly universally highly expressed by neuroblastomas regardless of disease stage [33, 34]. The SK-N-BE(2) cell line, derived from a patient with relapsed, high-risk neuroblastoma, has been well-established as a clinically relevant model of high-risk disease [35, 36] given its MYCN amplification and p53 mutation conferring chemoresistance [37]. As its GD2 expression was quantified previously by our group [21] prior to selection for xenograft generation, SK-N-BE(2) does provide a well-characterized platform for evaluating a GD2-targeted imaging agent. Secondly, while a formal receptor blocking study with excess unlabeled Dinutuximab was not performed, we were able to demonstrate specificity of the tracer for neuroblastoma through comparison to an isotype-control tracer, reiterating this specificity from both optical and gamma standpoints. GD2 binding of ^111^In-Dinutuximab-IRDye800 was preserved as demonstrated by ELISA assay. Lastly, formal pharmacokinetic modeling and radiation dosimetry analyses were beyond the scope of the present optimization study. While biodistribution studies identified four days after administration as the optimal imaging window, future studies will incorporate pharmacokinetic, dosimetric, and safety assessments to further characterize the translational and clinical profile of the tracer.

## Conclusion

Here, we demonstrate with ^111^In-Dinutuximab-IRDye800 that a dual-modality, GD2-targeted intraoperative imaging is achievable within a well-defined and clinically translatable formulation. The systematic optimization described yielded an agent with preserved targeting specificity, adequate fluorescent and radioactive signal, and favorable *in vivo* tolerability. In a surgical model designed to reflect clinical trial conditions, the tracer was able to detect lesions not seen under normal white-light conditions, thus demonstrating the potential to positively aid intraoperative decision-making. For a disease in which the completeness of surgical resection directly affects outcome and complication rates are unacceptably high, these results provide the foundation for advancing ^111^In-Dinutuximab-IRDye800 towards further Investigational New Drug (IND)-enabling studies and ultimately first-in-human evaluation.

## Abbreviations

EFS: event-free survival; IMI: intraoperative molecular imaging; FGS: fluorescence-guided surgery; RGS: radio-guided surgery; ICG: indocyanine green; EPR: enhanced permeability and retention; EGFR: epidermal growth factor receptor; CEA: carcinoembryonic antigen; DTPA: diethylenetriaminepentaacetic acid; NIR: near-infrared; ROI: regions of interest; MFI: mean fluorescence intensity; CSE: clinically significant events; PPV: positive predictive value; NPV: negative predictive value

## Acknowledgements

This work utilized the Hillman Cancer Center Small Animal Multimodal Imaging and Radiopharmaceuticals Application, a shared resource at the University of Pittsburgh supported by the CCSG P30 CA047904.

## Ethics Committee Approval and Patient Consent

Animal studies were performed in accordance with the protocols approved by the Institutional Animal Care and Use Committee of the University of Pittsburgh (IACUC protocol number: 24013900)

